# Historical overestimated abundance of a Critically Endangered species: the case of the tamaraw (*Bubalus mindorensis*) of Mindoro Island

**DOI:** 10.64898/2026.11.13.766428

**Authors:** Alexis Rutschmann, Andrew Tilker, Fernando García-Gil, Emmanuel Schütz, Christophe Bonenfant

**Affiliations:** Sorbonne Université, UPEC, CNRS, IRD, INRA, Institut d’Écologie et des Sciences de l’Environnement, IEES, Paris, France; Re:wild, Austin, Texas USA; Leibniz Institute for Zoo and Wildlife Research, Berlin, Germany; D’ABOVILLE Foundation and Demo Farm Inc, Puerto Galera, Philippines; Université de Lyon, F-69 000, Lyon; Université Lyon 1; CNRS, UMR 5558, Laboratoire de Biométrie et Biologie Évolutive, F-69 622, Villeurbanne, France

**Keywords:** _abundance_, *_Bubalus mindorensis_*, _conservation planning_, _density_, _detection probability_, _independent double-observer method_, _population trend_

## Abstract

Population abundance is a central parameter in conservation, for which biased estimates can seriously undermine management strategies for threatened species. The tamaraw (*Bubalus mindorensis*) is a Critically Endangered large herbivore endemic to Mindoro Island, Philippines. The species has been monitored annually at Mounts Iglit-Baco Natural Park (MIBNP) since 2000 using a multi-vantage-point count method that is prone to both imperfect detection and repeated counting of the same individuals across sessions. Because the historical abundance estimation is an index of relative abundance with no probabilistic modeling of counts, the actual number of tamaraw in what is considered as the largest remaining population is still poorly known. To fill this gap, we conducted an independent double-observer experiment over five years (2022–2026) and fitted Bayesian hierarchical detection models to the counts. As expected, detection probability was < 1 throughout (single-observer estimates 0.41–0.84; combined-observer 0.68–0.96), confirming that a substantial fraction of animals is missed by rangers during all counts. Compared with these estimates, historical counts overestimated tamaraw abundance by a factor of 1.61–2.34, yielding a mean population size across 2022–2026 of 203 individuals (95% credible interval 182–226) rather than the roughly 400–500 individuals previously assumed. Our results imply that the MIBNP tamaraw population, while more vulnerable to demographic and genetic risks than previously believed, has likely responded positively to ongoing conservation efforts; we recommend that population viability assessments and translocation plans for the species be revised accordingly.

## INTRODUCTION

Information about population abundance (*i.e.*, number of animals, or their density in a given area) is critical to understanding the past, present, and future trajectory of species in a changing environment (Caughley 1977; Callaghan et al. 2024). In particular, robust estimates of population abundance are crucial to endangered species conservation, since strategies and management policies are often based on the number of animals left in the wild (Mace et al. 2008; Neel et al. 2012). Furthermore, many conservation frameworks use population size, in conjunction with other criteria, to assess species status (*e.g.* IUCN, NatureServe, UE Bird Directive). For instance, according to the IUCN Red List Categories and Criteria, a species should be classified critically endangered if, among other things, its population size is <250 individuals (IUCN 2012, p. 20). Unfortunately, accurately estimating abundance of wild populations is, in practice, still a major challenge (Callaghan et al. 2024).

The main difficulty in assessing the abundance of a population is the imperfect detection of wildlife during surveys (Andersen 1953; Anderson 2001). In most situations, an observer will miss one or more individuals during a survey, for example when visibility is poor (*e.g.*, dense vegetation cover), animals are elusive, (*e.g.*, color, size, cryptic behavior, avoidance of humans MacKenzie et al. 2005), or individuals cannot be reliably identified (Johansson et al. 2020). The experience of the observer can also play a role in detection rates (Garel et al. 2005; Sunde & Jessen 2013). Because the number and proportion of animals missed in these situations is unknown, uncorrected counts generally result in an underestimation of abundance (*i.e.* negative bias Andersen 1953). Such an underestimation has the potential to mislead population management decisions (Kéry & Schmidt 2008). For example, in western Europe, the consistent underestimation of roe deer (*Capreolus capreolus*), red deer (*Cervus elaphus*) and wild boar (*Sus scrofa*) densities has led to conservative harvest quotas and, as a result, “overabundant” populations in some areas (Apollonio et al. 2010).

The classic way to deal with an underestimation of animal abundance is to estimate the detection probability of individuals to infer population abundance. Statistical methods designed to do this have been under active and continuous development since the seminal publication of Petersen (1896). Since then, two main methodological approaches have prevailed: capture-mark-recapture (CMR, Otis et al. 1978) and distance sampling (DS, Buckland et al. 2004; Burnham et al. 1980). These two estimators of population abundance have since inspired the development of many other approaches (Seber 1982; Schwarz & Seber 1999) with the primary goal of estimating a detection probability and correcting for the negative bias of raw counts. More recently, spatially replicated counts (*N*-mixture models, Royle 2004) and random-encounter diffusion models (Rowcliffe et al. 2008) have emerged as new credible alternatives to CMR and DS to measure population abundance from unmarked animals in the wild (but see Barker et al. 2018, for discussion).

While the underestimation of animal abundance may be seen as conservative for the conservation of a species, its overestimation may have significant and more concerning implications. Although much less frequently reported than underestimation cases, overestimation of population abundance can lead to overly optimistic estimates of extinction risk (see Garshelis et al. 2008, for an example on Giant panda (*Ailuropoda melanoleuca*)). Contrasting situations leading to an overestimation of abundance have been reported in the literature when applying the usual estimators of population abundance (Table 1). For example, when using a noninvasive capture-mark-recapture estimator of abundance based on hair or feces collections, genotyping errors are known to lead to an overestimation when not accounted for because it leads to an apparent larger number of different individuals (Lukacs & Burnham 2005). Similarly, population abundance overestimation may occur when the underlying hypotheses of the estimators are not met, for example when observers fail to detect animals located right on the transects during distance sampling (see Turnock & Quinn 1991, when detection distance is zero). Overestimation of abundance can also result from a poorly defined sampling design (Oberosler et al. 2022; Oyster et al. 2018), or from a poor assessment of the geographic range or activity of animals when converting a population size into a density estimate (Cheyne et al. 2013; Miller et al. 2005). Finally, when animals are counted more than once during a survey, there may also be an overestimation of population size (see Linklater & Cameron 2002; Bonenfant et al. 2023b, for examples on feral horses (*Equus caballus*) and giraffes (*Giraffa camelopardalis*)).

**Table 1.** Estimators of population abundance that accommodate for multiple observations of the same individuals in animal counts. Multiple observations of the same animals during counts may lead to strong overestimation of population size if not accounted for. Abundance overestimation is particularly dangerous in conservation biology because fewer animals than anticipated may be left in the wild.

| Estimator | Principle | Reference |
|---|---|---|
| *N*-mixture | Fitting a Poisson-binomial mixture model to estimate a detection probability *p* | Kéry & Royle (2020) |
| mark-resight | Frequency distribution of marked animals is applied to unmarked animals | Arnason et al. (1991) |
| robust design (temporary emigration) | Explicitly models whether a marked individual is present/available at a site on a given occasion as a latent state, distinguishing genuine absence from unavailability due to movement, rather than treating repeat captures of a moving individual as independent | Kendall et al. (1997) |
| *N*-mixture with temporary emigration | Extends the closed-population *N*-mixture model with an explicit availability/movement parameter for unmarked individuals, relaxing the closure assumption that would otherwise let the same moving individual be recounted across sites or occasions | Chandler et al. (2011) |

The tamaraw (*Bubalus mindorensis*) is a wild cattle species endemic to the island of Mindoro in the Philippines (Custodio et al. 1996). The tamaraw once occurred in large numbers across the island, but its numbers have drastically declined over the last century to less than 500 animals that occur in three isolated populations (Long et al. 2018). The largest population occurs in Mounts Iglit-Baco Natural Park (MIBNP), located in uplands, Western coast of Mindoro (Long et al. 2018; Matsubayashi et al. 2010). Since 2000, annual counts of tamaraw have been carried out with the aim of monitoring annual population size and assessing the effectiveness of conservation measures and actions. Conservation attention in the past 30 years has led to an apparent growth in the MIBNP population (Fig. 1, Bonenfant et al. 2023a). Tamaraw counts are conducted visually, with observers stationed on vantage points within the core population area, with some animals missed during the counting sessions. At the same time, counts are repeated and cumulated over eight consecutive counting sessions. With such a sampling design, multiple counts of the same individuals are highly likely, albeit to an unknown extent. The annual counting data of tamaraw in MIBNP is unique in that it combines the usual underestimation issues of raw counts (*i.e.* imperfect detection) with major risks of overestimation (*i.e.* multiple counts between and within sessions).

**Figure 1.**
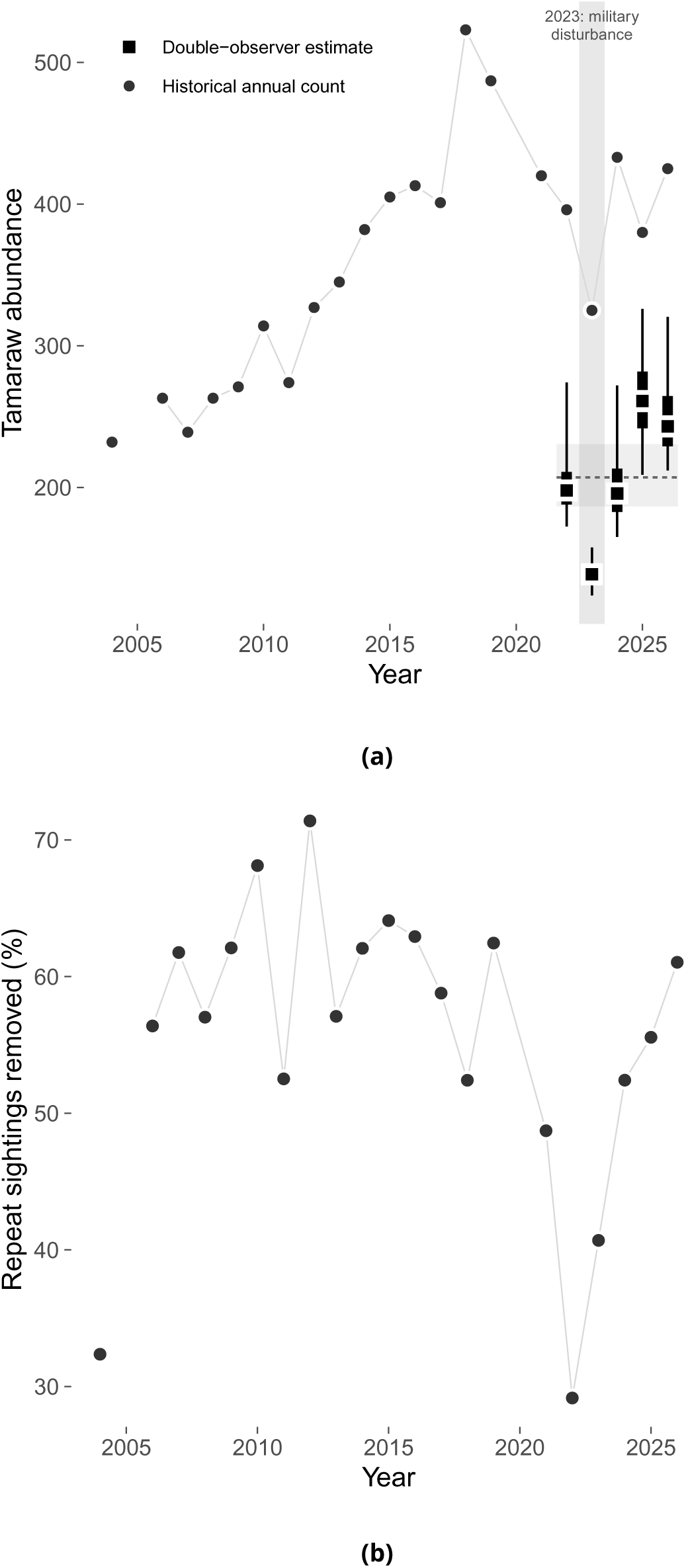
Historical results of the annual tamaraw population count, compared to the double-observer estimate. *a.* The total number of animals estimated after consolidation each year (circles, connected by a light guide line), together with the independent double-observer abundance estimate (squares) for the five surveyed years – illustrating the large, consistent gap between the two methods. Double-observer estimates are shown as point-intervals: the square marks the posterior median, the thick bar the 50% credible interval, and the thin bar the 95% credible interval. The dashed line and shaded band show the state-space model’s mean abundance and 95% credible interval over 2022–2026, with year-specific sampling variance removed. *b.* The yearly variation in the proportion of tamaraw number removed from the raw annual counts because they are considered as previously observed. Note that there was no count in 2020 due to the COVID-19 pandemic.

Although the annual counts have been instrumental in building a time series of tamaraw abundance (Bonenfant et al. 2023a), this historical method has never been critically evaluated and compared to a proper population size estimator accounting for the detection probability of animals in the field (see Douhard et al. 2013; Garel et al. 2005; Pellerin et al. 2017, for examples of empirical validation of population indices on large herbivores in Europe). Despite more than 20 years of annual counting data, conservation scientists are therefore faced with a situation in which the actual number of tamaraw within MIBNP is still largely unknown. Given the tamaraw’s Critically Endangered status (Boyles et al. 2016), uncertainty in the tamaraw population status in MIBNP is problematic because it hinders effective conservation planning and robust monitoring of the single largest population of the species. Here, we attempt to provide the first proper estimate of the MIBNP tamaraw population by implementing an independent double-observer experiment (Nichols et al. 2000) over five survey years (2022, 2023, 2024, 2025 and 2026) that explicitly accounts for the imperfect detection of tamaraw by observers.

## MATERIAL AND METHODS

### Study area

Mounts Iglit-Baco Natural Park (N12^◦^54^′^, E121^◦^13^′^) is a 106 655 ha protected area in the south-central part of Mindoro Island (Philippines). Within the park, a core zone of monitoring (CZM, approximately 3 000 ha, about 3% of MIBNP) is patrolled on a regular basis by park rangers, with limited potential for range expansion (Bonenfant et al. 2023a). Within the CZM, a smaller no-hunting zone established in 2016 (approximately 1 600 ha, about 1.5% of MIBNP) partly benefits from a non-hunting agreement with local indigenous people living inside the park (Bonenfant et al. 2023a). The CZM area is a rolling grassland plateau with an average elevation of 800 m a.s.l., dominated by cogon (*Imperata cylindrica*) and wild sugarcane (*Saccharum spontaneum*), and interspersed with numerous wooded creeks, secondary forest fragments, and steep hills. The tropical climate is strongly seasonal with a rainy season ranging from June to October (mean temperature 26.5°C; mean rainfall: 150 mm), and a hot dry season ranging from November to May (mean temperature 29.8°C; mean rainfall: 5 mm). Apart from humans, the only potential predator of tamaraw is the reticulated python (*Malayopython reticulatus*) that is capable of preying on calves or yearlings, although no such predation observations have been made to date.

The CZM area hosts the largest remaining tamaraw population. Although unfenced, we assumed that the population would remain demographically closed during sampling. This assumption is likely to be met because the natural mortality risk for this long-lived species is assumed to be low (Gaillard et al. 2000). We know that some individuals are harvested by indigenous people near the periphery of the no-hunting area, but traditional hunting generally occurs in months of June to November. The recruitment of newborns in a two-week period is also unlikely, because even though the timing of the births of female tamaraw in MIBNP is not known in detail, rangers have never reported observations of newborn or very young tamaraw during sampling. Similarly, there are no known populations of tamaraw near the CZM (Long et al. 2018), and thus it is not possible that immigrating animals would have entered from outside the study during survey.

### Historical counts of tamaraw

Since 2000, the Tamaraw Conservation Program Office (TCPO), an office of the Philippine Department of Environment and Natural Resources, has been organizing annual tamaraw counts throughout the CZM. Rangers and volunteers are dispatched across 19 vantage points, covering nearly 2 200 ha within the CZM (Fig. 1). For four consecutive days, tamaraw are counted simultaneously twice a day at dusk and dawn, with each session lasting 1.5 hours. At each vantage point, at least two observers were used to record every animal observed, along with observation time, location, sex, and age-category (calf, yearling, subadult, or adult).

At the end of each session, group composition, position, sex, age and direction of movement were used by observers to attempt to identify animals seen simultaneously from adjacent vantage points or in previous sessions. Based on this information, observers could decide which tamaraw groups were to be removed from the total count to yield a so-called *consolidated* population size of tamaraw. This *consolidated* estimate is still the official abundance estimate of the tamaraw population for the Philippine authorities to set conservation actions for the species (see Bonenfant et al. 2023a, for more details). We denote this official, park-wide consolidated count for year *t* as *C_t_*. An important point is that there is no attempt to apply any population size estimator to these count data. In other words, the official tamaraw abundance is a measure of relative abundance, not accounting for the imperfect detection of animals, although attempting to reduce potential multiple counts of animals.

### The double-observer methodology

To obtain the first robust estimate of the size of the tamaraw population in MIBNP, we used the independent version of the double-observer estimator of population size (Nichols et al. 2000) as follows. In April 2022, May 2023, April 2024, May 2025 and April 2026, after annual population counts, five teams of two people (denoted *A* and *B*) concurrently and independently recorded animals at 15–18 vantage points (VPs) depending on the year (15 in 2022; 18 in 2023, 2024, 2025 and 2026, with the addition of the Iyan, Malibayong and Tangle VPs). Due to limited manpower and logistic resources, we excluded VPs having very low numbers of individuals recorded during previous counts (< 2 animals/year). We assigned VPs to observer teams in time and space to minimize the chances of multiple observations of the same animal.

Observations spanned two periods of 15 minutes each, separated by a 45-minute break time during which observers refrained from observing the landscape. We conducted 2 × 15 minutes of observation sessions, four times over two days, at each of the *k* ∈ {15, 18} vantage points, for a total of *n* ∈ {121, 181, 144, 144, 144} site × date × session replicates for years 2022, 2023, 2024, 2025 and 2026 (indexed by *t*) respectively. Because the double-observer model assumes independence of counts, every effort was made to keep observers independent, including that they did not communicate with one another and stayed at least two meters apart at all times. Immediately after the 15-minute sessions ended, the two observers *A* and *B* compared their results to get the number of tamaraw seen by *A* only 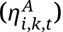, by *B* only 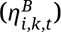, and seen by both *A* and *B* 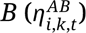. Those numbers were further split into different sex-(male and female) and age-categories because body size can influence detection probability of animals (McCorquodale 2001). Note that age-class definitions differed slightly between years.

#### Statistical analyses

We implemented the independent double-observer estimator (Nichols et al. 2000) in a Bayesian framework (Roy et al. 2022; Wilson et al. 2014). We used the Poisson-multinomial equivalence of the model likelihood, which avoids having to treat the number of individuals missed by both observers as an explicit latent variable (Baker 1994; Royle 2004). For each VP × date × sex × age-class combination (row *i*), the number of tamaraw seen by *A* only, by *B* only, or by both were conditionally independent Poisson counts with mean *N_i_* × *π_i_*, *_j_*, where *j* = 1 (seen by *A* only), *j* = 2 (seen by *B* only) and *j* = 3 (seen by both), and *N_i_* is the expected local abundance for the corresponding combination of time, location and tamaraw category. Writing 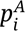and 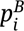for the row-specific detection probabilities of observers *A* and *B* (obtained from the same regression below, evaluated at the observer-role level of ***X****_p_*_,*i*_ corresponding to each observer), the double-observer probabilities follow from the assumption of independent detection:

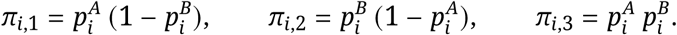

We modeled 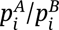 and *N_i_* with fixed effects and VP-level random intercepts on the logit and log scale respectively:

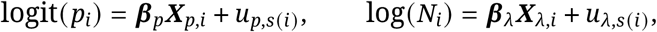

where ***β****_p_* and ***β****_λ_* are the vectors of fixed-effect coefficients associated with the design matrices ***X****_p_*_,*i*_ and ***X****_λ_*_,*i*_; *s*(*i*) ∈ {1, …, *k*} is the VP of row *i*; ***X****_p_*_,*i*_ included weather (good/poor) and observer role (*A*/*B*); ***X****_λ_*_,*i*_ included sex and age class of tamaraw (reproductive status in 2022–2024, four age classes in 2026). The VP random intercepts 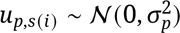 and 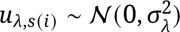 capture, respectively, consistent differences in detectability and spatial variation in local density among VPs not accounted for by the fixed effects, with *σ_p_* and *σ_λ_* their standard deviations across VPs. We also tested whether replacing *u_λ_*_,*s*(_*_i_*_)_ with a spatially explicit structure of residuals (conditional autoregressive or CAR) improved the fit (Supplementary material, S2). CAR models did not perform better than the fully independent model, so we kept the simpler unstructured version reported here.

For each year, we calculated the park-wide population size *N̂_t_* by averaging *N_i_* across repeat survey days within each VP × sex × age-class group, and summing across groups and VPs, within each posterior draw. The same procedure restricted to rows at a single VP *s*, summed across sex/age groups but not across VPs, gives the VP-level abundance estimate *N̂_s_* reported in Table 4. We obtained point estimates and their precision by taking the median and 95% credible interval. For the three VPs not yet sampled in 2022 (Iyan, Malibayong, Tangle), *N_i_* was imputed using a historically-calibrated prior. We regressed the VP-level random effect fitted at the 15 surveyed VPs (*u_λ_*_,*s*(_*_i_*_)_) against log-transformed historical counts (2004–2019) at those same VPs (*C_t_*_,*i*_), and the fitted relationship informed the missing VPs’ posteriors. Our model assumed demographic closure within each sampling period (2 days overall), a fair assumption given the tamaraw’s low natural mortality in a 2-day time window, and the no-hunting protection area covering much of our study site (Bonenfant et al. 2023b).

The five annual estimates of tamaraw population size *N̂_t_* were then analyzed with a hierarchical Bayesian state-space model to tease apart sampling from process variances (Kéry & Royle 2020). We approximated the posterior distribution of *N̂_t_* by its mean and standard deviation on the log scale (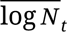 and *s_t_*). We treated 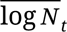 as the realization of a normal distribution whose parameters are the mean of the “true” population size 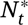 (hereafter the *state-space estimate* once fitted, denoted 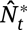) and the sampling variance 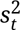:

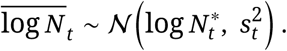

An exponential demographic model linked log 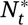 with log 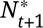, with a separate growth rate *r_t_* for every year-to-year transition, equivalent to a random walk on the log scale:

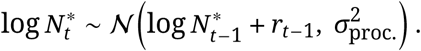

Where 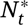 are the annual estimates of the tamaraw population size at Mounts Iglit-Baco with sampling variance removed. Annual growth rates *r_t_* were drawn from a common hierarchical distribution, 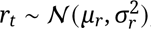, with *μ_r_* the mean annual growth rate – denoted *r̄* throughout this manuscript – and *σ_r_* its standard deviation across transitions. With no constraint on the prior distribution of *r_t_*, this model could return biologically implausible growth rates for the tamaraw. Published maximum growth rates for similarly-sized bovids under the most favorable ecological conditions rarely exceed ∼25–30% per year (Caughley 1977; Hone et al. 2010). We therefore truncated *r_t_* at log(1.3), setting a maximum annual growth rate (exp(*r_t_*) − 1) at +30%. This truncation was applied one-sided only, rate of decline staying unbounded. From the fitted state-space model, we report on three quantities: the mean annual growth rate (*r̄* ≡ *μ_r_*), the four annual growth rates *r_t_*, and the mean of 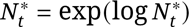 across all five years of study.

We fit the models with JAGS (Plummer et al. 2003) in the **R** environment (R Core Team 2025) with uninformative priors on all parameters other than for upper limit of annual growth rate. We ran 4 MCMC chains with 300 000 iteration, retaining 40 000 values to compute posterior distribution of estimated parameters, and checked convergence with the Gelman-Rubin *R̂* statistic (threshold < 1.1) and effective sample size (Brooks & Gelman 1998). We also checked the posterior-predictive (Bayesian *p*-value) goodness-of-fit. We report parameter estimates as posterior medians with 95% credible intervals (Louis & Zeger 2008): _95%_*_low_ estimate* _95%_*_up_*. All JAGS/R code, including a fully self-contained, reproducible worked example, is freely available on GitLab at: https://gitlab.in2p3.fr/christophe.bonenfant/tamaraw-abundance.

## RESULTS

### Temporal variation of historical tamaraw counts at MIBNP

The historical annual count time series available to us begins in 2004 and shows tamaraw abundance increasing markedly within the MIBNP, from *C*_2004_ = 232 individuals in 2004 to a maximum of *C*_2018_ = 523 in 2018. After that, tamaraw numbers have fluctuated without a clear further increase: *C*_2019_ = 487 in 2019, *C*_2021_ = 420 in 2021, *C*_2022_ = 396 in 2022, *C*_2023_ = 325 in 2023, *C*_2024_ = 433 in 2024, *C*_2025_ = 380 in 2025 and *C*_2026_ = 425 in 2026 (Fig. 1). The mean population size of tamaraw according to the multi-vantage-point count method was *C̄* = _320.7_ 358.0 _395.3_ individuals over 2004–2026. Restricted to 2022–2026, the period over which double-observer surveys were also conducted (below), the mean historical count was *C̄* = 391.8 individuals. On average the consolidated numbers, that is the number of detected tamaraw after suspected multiple counts have been removed by experienced rangers, comprised 44.3% of the total number of seen animals (see Supplementary material, S1 for details).

### Tamaraw population size with double-observer estimator

We carried out *n* = 121, 181, 144, 144 and 144 vantage point × date × session replicates in 2022, 2023, 2024, 2025 and 2026 respectively for the double-observer experiment. We counted a total of 450, 369, 448, 456 and 611 tamaraw (raw, uncorrected for detection or multiple counts) across all replicates in 2022, 2023, 2024, 2025 and 2026 respectively.

Detection probability was well below one in every year, confirming imperfect detection of tamaraw by observers: *p̂_A_* = _0.68_ 0.75 _0.81_ and *p̂_B_* = _0.66_ 0.73 _0.78_ in 2022; *p̂_A_* = _0.59_ 0.67 _0.74_ and *p̂_B_* = _0.68_ 0.77 _0.84_ in 2023; *p̂_A_* = _0.47_ 0.54 _0.68_ and *p̂_B_* = _0.48_ 0.56 _0.70_ in 2024; *p̂_A_* = _0.41_ 0.48 _0.58_ and *p̂_B_* = _0.45_ 0.54 _0.64_ in 2025; and *p̂_A_* = _0.54_ 0.60 _0.74_ and *p̂_B_* = _0.58_ 0.64 _0.78_ in 2026. Combining both observers, the overall detection probability by at least one observer was _0.89_ 0.93 _0.96_ (2022), _0.87_ 0.93 _0.96_ (2023), _0.73_ 0.80 _0.90_ (2024), _0.68_ 0.76 _0.85_ (2025) and _0.81_ 0.86 _0.94_ (2026) (Fig. 2). Detection probability also varied substantially among VPs for a given year (standard deviation of the VP-level random effect ranging from *σ̂_p_* = 0.11 to *σ̂_p_* = 0.31 across years and observers) suggestive marked spatial heterogeneity in detectability of tamaraw between VPs.

**Figure 2.**
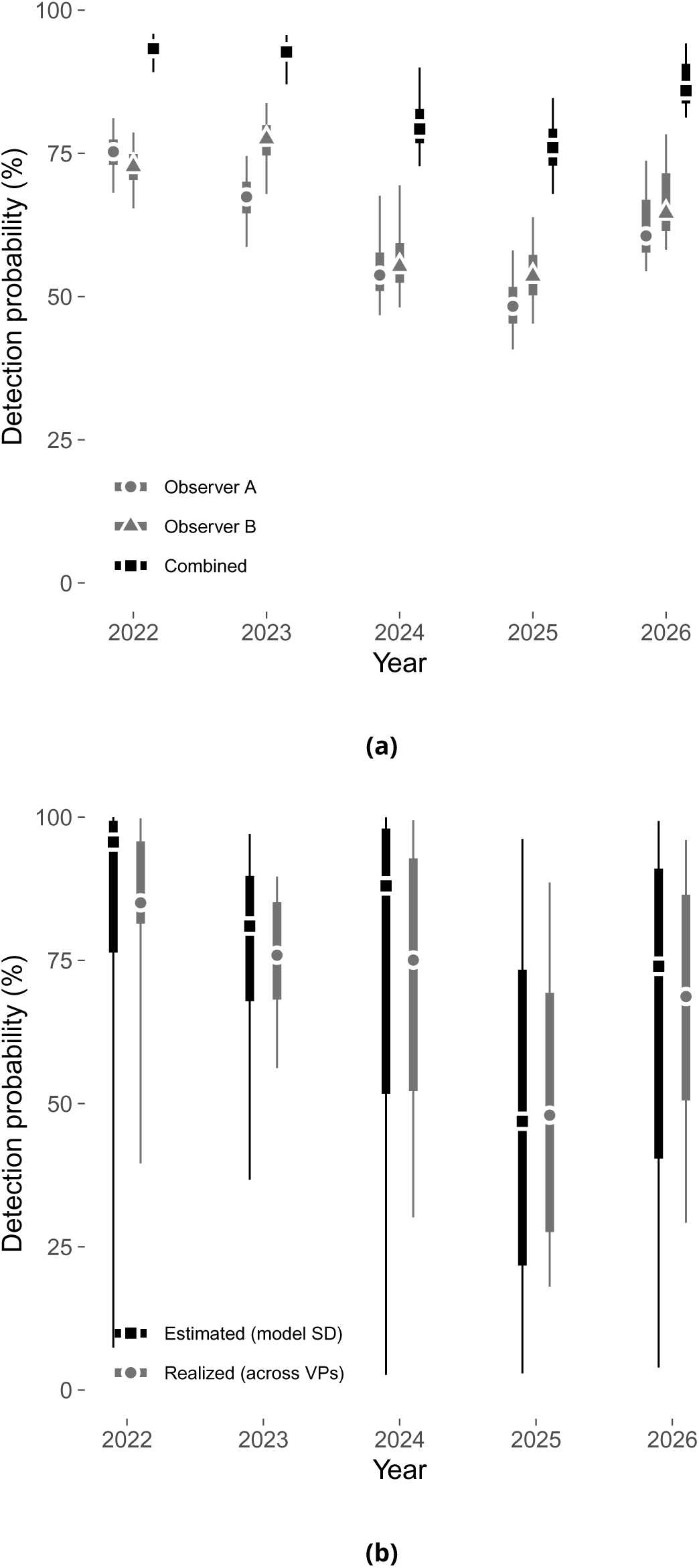
Detection probability of tamaraw by observer and vantage point, 2022–2026. *a.* Estimated detection probability of Observer A, Observer B, and the two observers combined, by year, shown as point-intervals: the point marks the posterior median, the thick bar the 50% credible interval, and the thin bar the 95% credible interval. *b.* Spread of detection probability among vantage points (VP), by year: the *Estimated* series is the VP-to-VP spread as returned by the fitted hierarchical random-effect standard deviation, and the *Realized* series is the actual spread of the site-level posterior means across that year’s surveyed vantage points. Same point-interval convention as *a*.

The total population size of tamaraw at MIBNP as determined by the double observer estimator was *N̂* = _172.5_ 197.9 _274.1_ in 2022, *N̂* = _123.7_ 138.7 _157.6_ in 2023, *N̂* = _165.1_ 195.7 _272.0_ in 2024, *N̂* = _198.4_ 236.1 _320.4_ in 2025, and *N̂* = _212.0_ 243.1 _320.4_ in 2026. In addition to time, the estimated number of tamaraw varied spatially to a large extent, with standard deviation of the VP-level random effect ranging from *σ̂_λ_* = 1.13 to *σ̂_λ_* = 1.44 across years. In 2026, the most recently and completely surveyed year, VP-level abundance ranged from *N̂_s_* = _31.3_ 38.4 _46.8_ at Bato Fidel to *N̂_s_* = _0.06_ 1.9 _28.1_ at Malibayong (Table 4). The correlation between fitted per-VP abundance and the raw (uncorrected) counts observed at the same VP *R*^2^ ranged from 0.845 to 0.994 across the five survey years.

Combining all five survey years in a hierarchical state-space model, tamaraw abundance at MIBNP increased over 2022–2026 at an estimated mean annual growth rate of *r̄* = _−21.6%_ +7.6% _+49.6%_ per year. The probability of a positive population growth over the study period was *P*(*r* > 0) = 0.69 (Fig. 1a, Fig. 3). Individual year-to-year transitions were more imprecisely estimated, with values of *r^*_2022_ = _−35.5%_ −3.2% _+27.4%_; *r^*_2023_ = _−25.1%_ +10.7% _+29.2%_; *r^*_2024_ = _−26.8%_ +8.5% _+28.9%_; *r̂*_2025_ = _−28.8%_ +5.5% _+28.5%_; (Fig. 3). With year-specific sampling variance removed, annual population sizes were 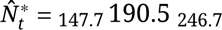 in 2022, _126.2_ 143.0 _162.2_ in 2023, _156.0_ 193.5 _245.2_ in 2024, _187.6_ 233.4 _293.5_ in 2025, and _201.0_ 246.8 _304.0_ in 2026, converging on a mean tamaraw population size at MIBNP of *N̂*^∗^ = _182.0_ 202.6 _226.1_ over 2022–2026.

**Figure 3.**
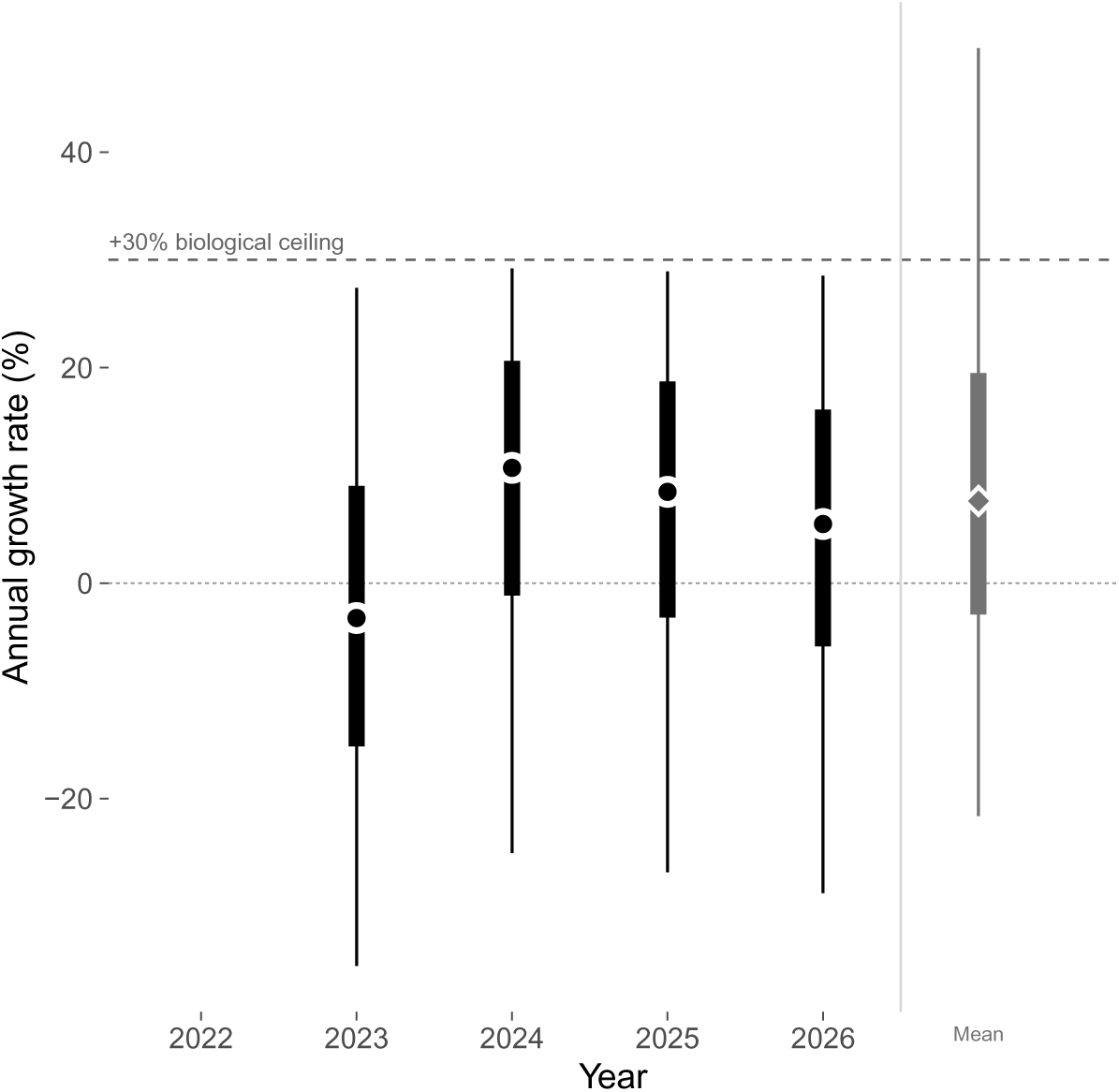
Annual population growth rate of tamaraw at MIBNP over time. Point-intervals of the state-space model’s annual growth rate, plotted at the end year of the counts (*e.g. r_t_* = log(*N_t_*/*N_t_*_+1_)). The point marks the posterior median, the thick bar the 50% credible interval, and the thin bar the 95% credible interval. On the right of the vertical divider, we also present the mean annual growth rate (*r̄*) over the full 2022–2026 period. The dashed horizontal line marks the +30% biological ceiling imposed on the growth-rate prior.

### Historical counts versus double-observer and state-space estimates

Table 3 compares, for each survey year, the historical (consolidated) count with our two model-based estimates of tamaraw abundance: the direct, single-year double-observer estimate *N̂*, and the state-space estimate *N̂* ^∗^ obtained once sampling variance is removed by pooling information across all five survey years. Both estimators of true abundance agree closely with one another (differences of 1–6 individuals per year) and are both consistently and substantially lower than the historical counts in every year. Historical counts overestimated tamaraw abundance relative to the double-observer estimate by a factor of 1.61 to 2.34 across years, and relative to the state-space estimate by a near-identical factor of 1.63 to 2.27. In both cases the smallest overestimation factor is found in 2025 and the largest in 2023, indicating that the historical overestimation of tamaraw abundance at MIBNP is a robust feature of the data, and not an artefact of treating each survey year in isolation from the others.

## DISCUSSION

Working with inaccurate population parameters has the potential to undermine conservation and wildlife management strategies (Anderson 2001; Nichols 2014). Our findings indicate that tamaraw abundance in Mounts Iglit-Baco Natural Park has been overestimated from historical counts over the last 6 years, and likely so for decades. Because Mounts Iglit-Baco Natural Park holds the largest population of tamaraw (Long et al. 2018), the overestimation of its population size puts the global conservation of the species in a more precarious light than previously anticipated, and will likely require stakeholders to re-evaluate management strategies in the protected area (DENR 2019).

### Biological insights from the double-observer methods

The historical tamaraw counts have been an activity of the Tamaraw Conservation Program Office (TCPO), ensuring continuous monitoring of the species since 2000. Our double-observer estimation of population size, however, suggests that actual tamaraw abundance is approximately half the population size values reported from the historical counts carried out at Mount Iglit-Baco Natural Park. Several lines of evidence support our interpretation of a long-lasting overestimation of tamaraw abundance in this protected area.

On biological grounds, the historical population size estimates appear implausible. Converting the historical population numbers to density (using the 2 200 ha counting area and the historical series’ maximum, *C*_2018_ = 523 in 2018) gives up to 24 individuals km^−2^. Such high numbers lie in the upper range of what is generally reported for other cattle species in their natural environment (see Table 2). This high population density of tamaraw is comparable to that reported for feral water buffalo (*Bubalus bubalis*) in Australia’s Northern Territory (up to 34 individuals per km²), where the species was introduced and has reached ecologically unsustainable levels, now recognized as a major environmental threat (Werner 2013). For comparison’s sake, population density of the anoa (*Bubalus depressicornis*), the closest relative of the tamaraw, was around 1 individual km^−2^ (Burton et al. 2005).

**Table 2.**
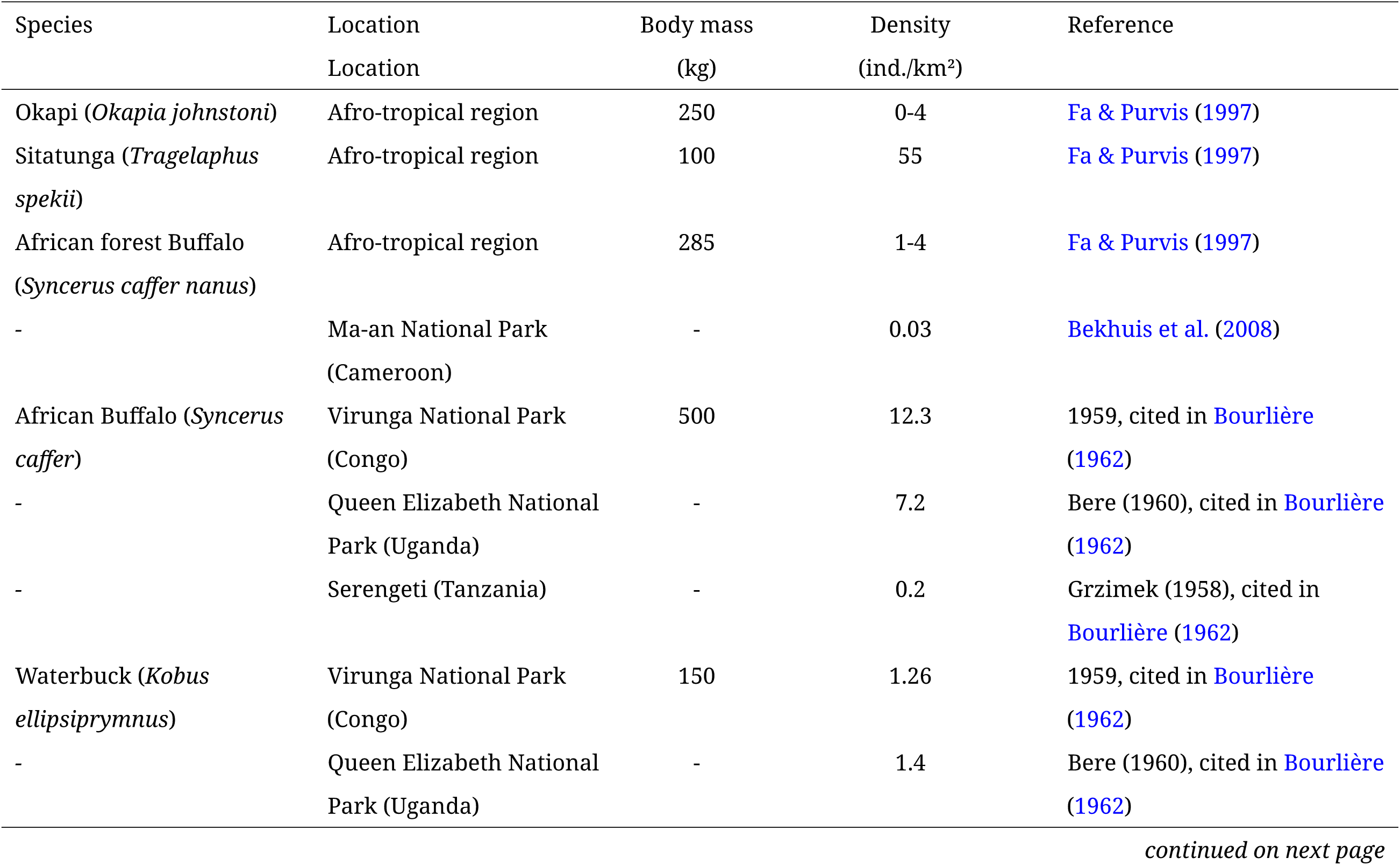
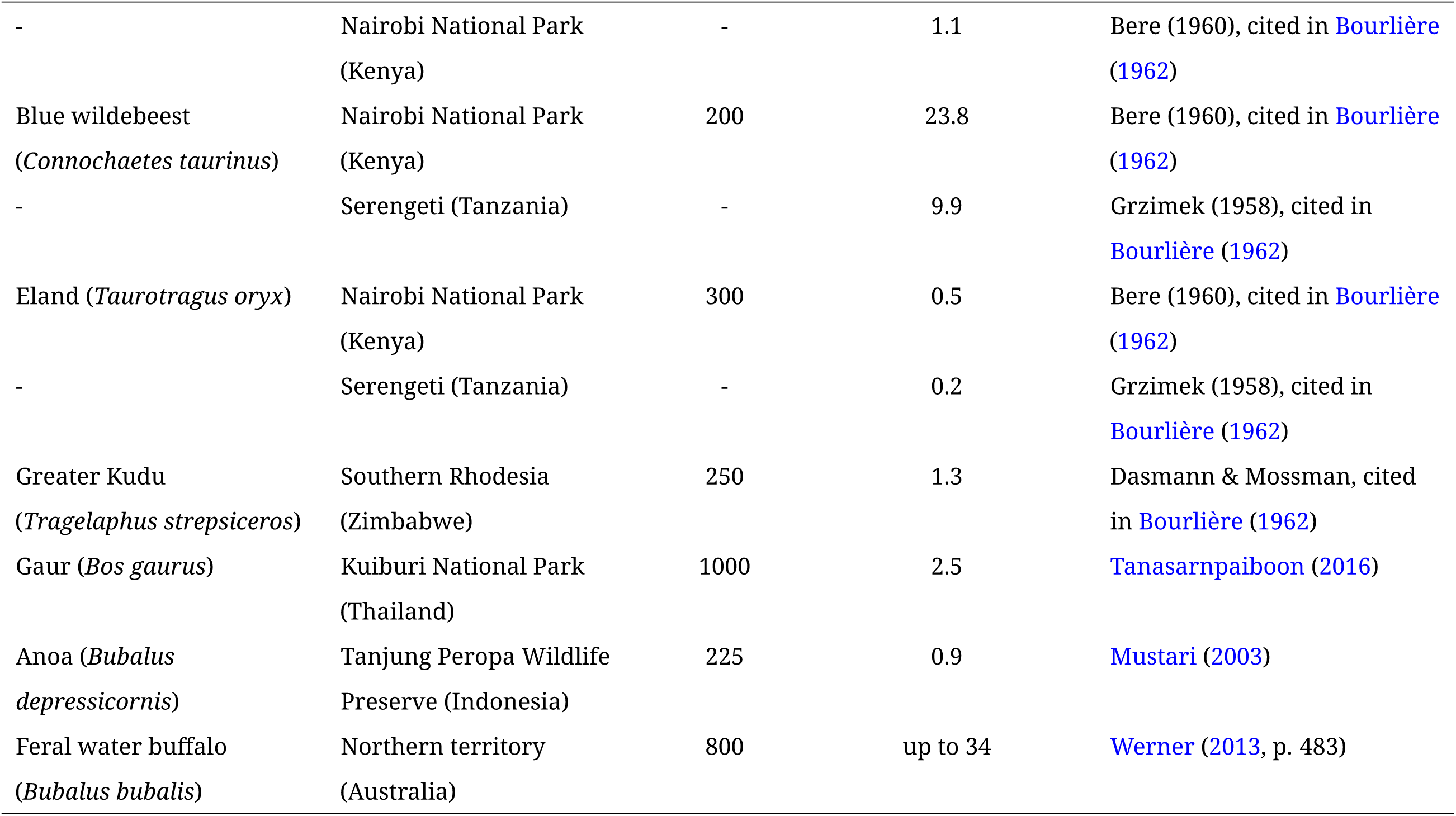
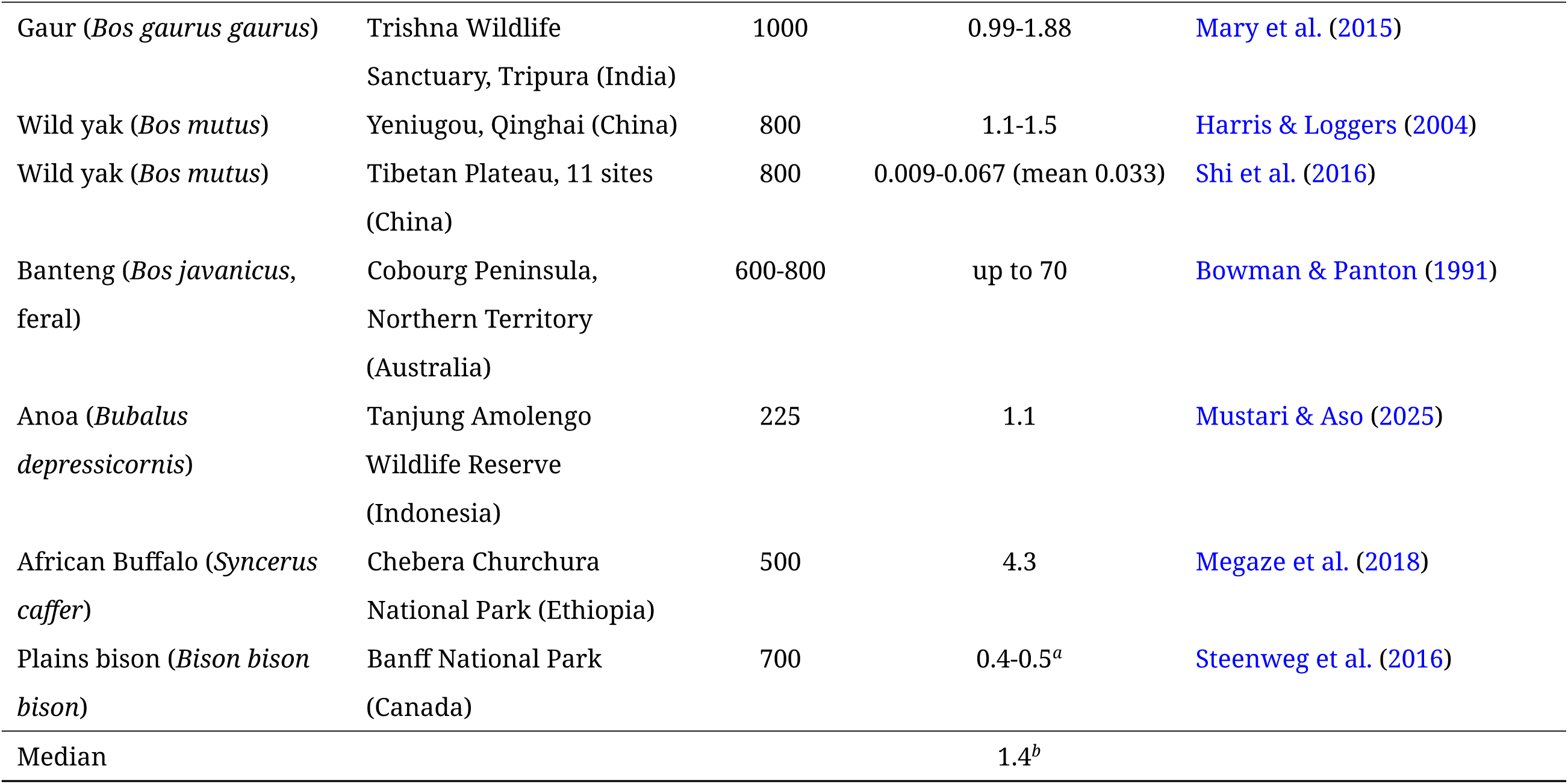
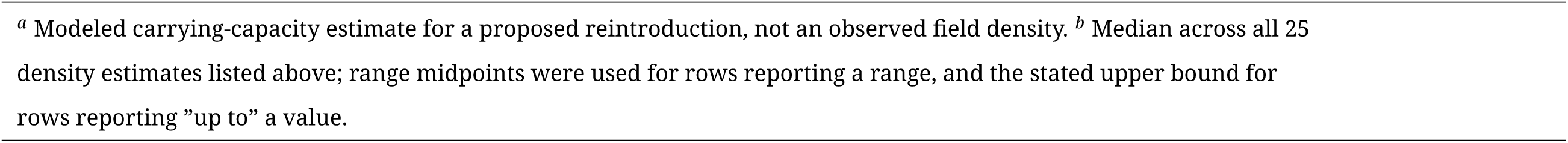
Estimates of population densities of several large Bovid species from a non-exhaustive review of the literature. The range of observed values suggests a rather high density of tamaraw (*Bubalus mindorensis*) on Mindoro Island, in the Mounts Iglit-Baco Natural Park.

| Species | Location | Body mass | Density | Reference |
|---|---|---|---|---|
|  | Location | (kg) | (ind./km<sup>2</sup>) |  |
| Okapi (*Okapia johnstoni*) | Afro-tropical region | 250 | 0-4 | Fa & Purvis (1997) |
| Sitatunga (*Tragelaphus spekii*) | Afro-tropical region | 100 | 55 | Fa & Purvis (1997) |
| African forest Buffalo (*Syncerus caffer nanus*) | Afro-tropical region | 285 | 1-4 | Fa & Purvis (1997) |
| - | Ma-an National Park (Cameroon) | - | 0.03 | Bekhuis et al. (2008) |
| African Buffalo (*Syncerus caffer*) | Virunga National Park (Congo) | 500 | 12.3 | 1959, cited in Bourlière (1962) |
| - | Queen Elizabeth National Park (Uganda) | - | 7.2 | Bere (1960), cited in Bourlière (1962) |
| - | Serengeti (Tanzania) | - | 0.2 | Grzimek (1958), cited in Bourlière (1962) |
| Waterbuck (*Kobus ellipsiprymnus*) | Virunga National Park (Congo) | 150 | 1.26 | 1959, cited in Bourlière (1962) |
| - | Queen Elizabeth National Park (Uganda) | - | 1.4 | Bere (1960), cited in Bourlière (1962) |
| - | Nairobi National Park (Kenya) | - | 1.1 | Bere (1960), cited in Bourlière (1962) |
| Blue wildebeest (*Connochaetes taurinus*) | Nairobi National Park (Kenya) | 200 | 23.8 | Bere (1960), cited in Bourlière (1962) |
| - | Serengeti (Tanzania) | - | 9.9 | Grzimek (1958), cited in Bourlière (1962) |
| Eland (*Taurotragus oryx*) | Nairobi National Park (Kenya) | 300 | 0.5 | Bere (1960), cited in Bourlière (1962) |
| - | Serengeti (Tanzania) | - | 0.2 | Grzimek (1958), cited in Bourlière (1962) |
| Greater Kudu (*Tragelaphus strepsiceros*) | Southern Rhodesia (Zimbabwe) | 250 | 1.3 | Dasmann & Mossman, cited in Bourlière (1962) |
| Gaur (*Bos gaurus*) | Kuiburi National Park (Thailand) | 1000 | 2.5 | Tanasarnpaiboon (2016) |
| Anoa (*Bubalus depressicornis*) | Tanjung Peropa Wildlife Preserve (Indonesia) | 225 | 0.9 | Mustari (2003) |
| Feral water buffalo (*Bubalus bubalis*) | Northern territory (Australia) | 800 | up to 34 | Werner (2013, p. 483) |
| Gaur (*Bos gaurus gaurus*) | Trishna Wildlife Sanctuary, Tripura (India) | 1000 | 0.99-1.88 | Mary et al. (2015) |
| Wild yak (*Bos mutus*) | Yeniugou, Qinghai (China) | 800 | 1.1-1.5 | Harris & Loggers (2004) |
| Wild yak (*Bos mutus*) | Tibetan Plateau, 11 sites (China) | 800 | 0.009-0.067 (mean 0.033) | Shi et al. (2016) |
| Banteng (*Bos javanicus*, feral) | Cobourg Peninsula, Northern Territory (Australia) | 600-800 | up to 70 | Bowman & Panton (1991) |
| Anoa (*Bubalus depressicornis*) | Tanjung Amolengo Wildlife Reserve (Indonesia) | 225 | 1.1 | Mustari & Aso (2025) |
| African Buffalo (*Syncerus caffer*) | Chebera Churchura National Park (Ethiopia) | 500 | 4.3 | Megaze et al. (2018) |
| Plains bison (*Bison bison bison*) | Banff National Park (Canada) | 700 | 0.4-0.5<sup>*a*</sup> | Steenweg et al. (2016) |
| Median |  |  | 1.4<sup>*b*</sup> |  |

<sup>*a*</sup> Modeled carrying-capacity estimate for a proposed reintroduction, not an observed field density. <sup>*b*</sup> Median across all 25 density estimates listed above; range midpoints were used for rows reporting a range, and the stated upper bound for rows reporting ”up to” a value.

**Table 3.**
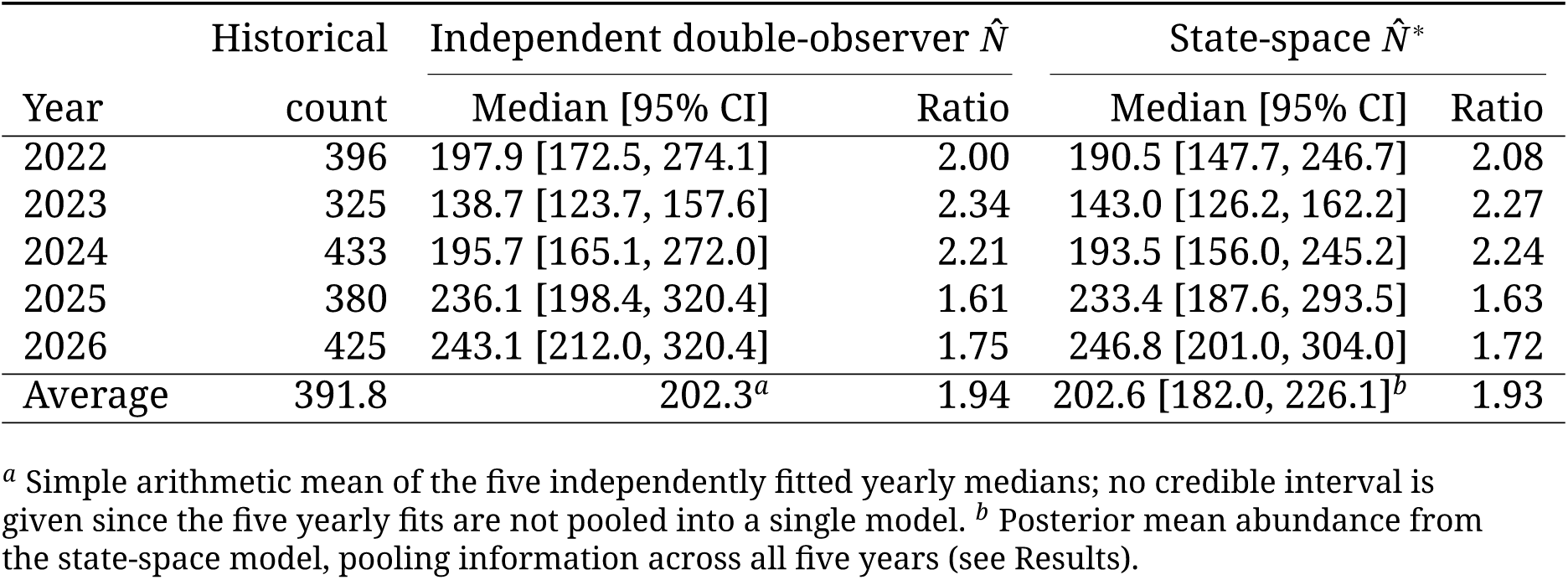
Comparison of the historical tamaraw count (official numbers released by the Department of Environment and Natural Resources of the Philippines) with the independent double-observer point estimate (*N̂*, fitted independently each year) and the state-space estimate (*N̂* ^∗^, sampling variance removed by pooling information across all five survey years) at Mounts Iglit-Baco Natural Park, 2022–2026. The two ratio columns tabulate the historical count divided by each model-based estimate for each year of the study.

|  | Historical | Independent double-observer *N̂* |  | State-space *N̂*<sup>\*</sup> |  |
|---|---|---|---|---|---|
| Year | count | Median [95% CI] | Ratio | Median [95% CI] | Ratio |
| 2022 | 396 | 197.9 [172.5, 274.1] | 2.00 | 190.5 [147.7, 246.7] | 2.08 |
| 2023 | 325 | 138.7 [123.7, 157.6] | 2.34 | 143.0 [126.2, 162.2] | 2.27 |
| 2024 | 433 | 195.7 [165.1, 272.0] | 2.21 | 193.5 [156.0, 245.2] | 2.24 |
| 2025 | 380 | 236.1 [198.4, 320.4] | 1.61 | 233.4 [187.6, 293.5] | 1.63 |
| 2026 | 425 | 243.1 [212.0, 320.4] | 1.75 | 246.8 [201.0, 304.0] | 1.72 |
| Average | 391.8 | 202.3<sup>*a*</sup> | 1.94 | 202.6 [182.0, 226.1]<sup>*b*</sup> | 1.93 |

<sup>*a*</sup> Simple arithmetic mean of the five independently fitted yearly medians; no credible interval is given since the five yearly fits are not pooled into a single model. <sup>*b*</sup> Posterior mean abundance from the state-space model, pooling information across all five years (see Results).

**Table 4.**
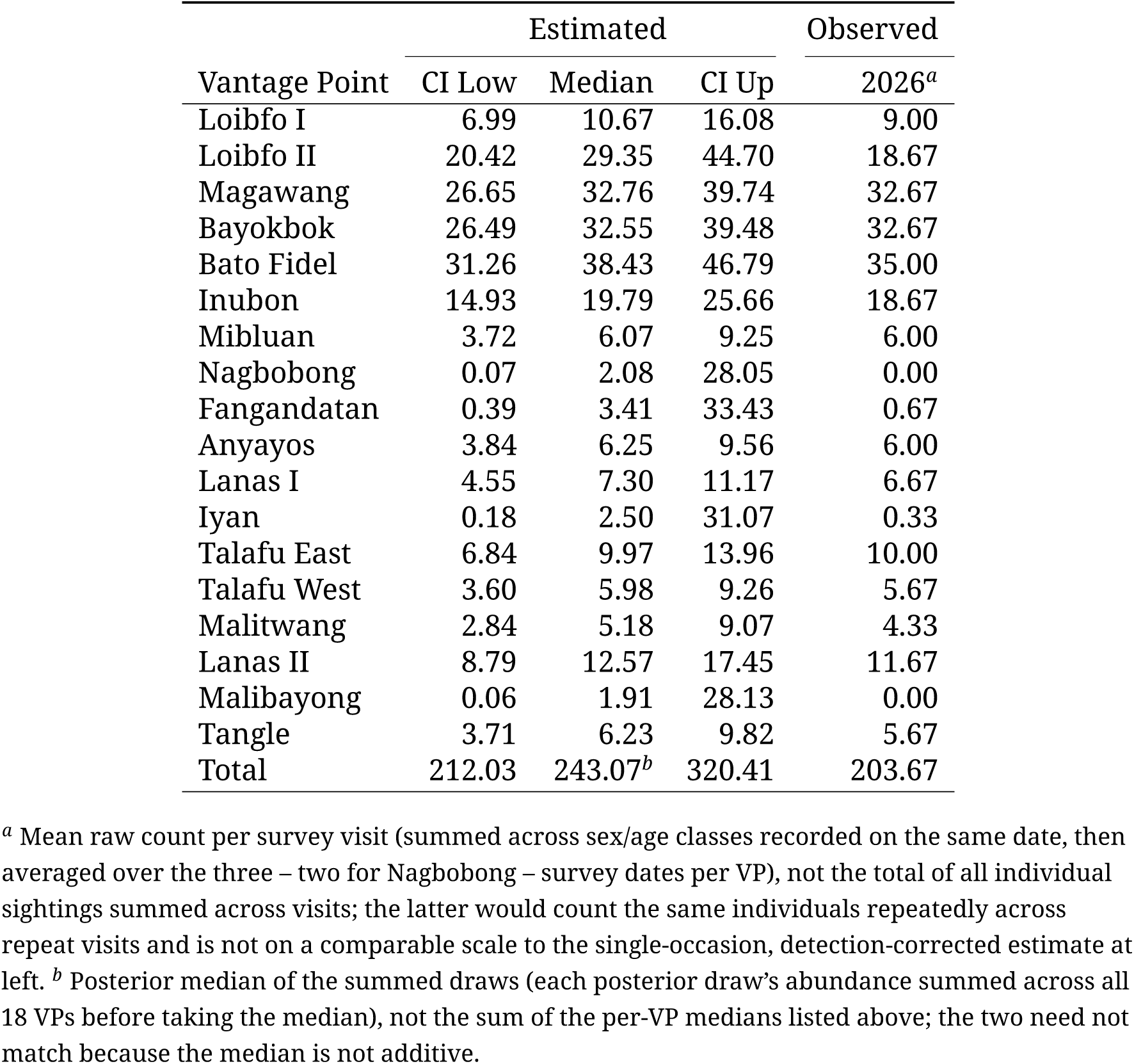
Spatial variation in population abundance of tamaraw (*Bubalus mindorensis*) on Mindoro Island, in the Mounts Iglit-Baco Natural Park, for the 2026 survey. We applied the independent double-observer estimator (Nichols et al. 2000) across the core zone of monitoring of this critically endangered species on a set of 18 vantage points distributed across the study site.

|  | Estimated |  |  | Observed |
|---|---|---|---|---|
| Vantage Point | CI Low | Median | CI Up | 2026<sup>*a*</sup> |
| Loibfo I | 6.99 | 10.67 | 16.08 | 9.00 |
| Loibfo II | 20.42 | 29.35 | 44.70 | 18.67 |
| Magawang | 26.65 | 32.76 | 39.74 | 32.67 |
| Bayokbok | 26.49 | 32.55 | 39.48 | 32.67 |
| Bato Fidel | 31.26 | 38.43 | 46.79 | 35.00 |
| Inubon | 14.93 | 19.79 | 25.66 | 18.67 |
| Mibluan | 3.72 | 6.07 | 9.25 | 6.00 |
| Nagbobong | 0.07 | 2.08 | 28.05 | 0.00 |
| Fangandatan | 0.39 | 3.41 | 33.43 | 0.67 |
| Anyayos | 3.84 | 6.25 | 9.56 | 6.00 |
| Lanas I | 4.55 | 7.30 | 11.17 | 6.67 |
| Iyan | 0.18 | 2.50 | 31.07 | 0.33 |
| Talafu East | 6.84 | 9.97 | 13.96 | 10.00 |
| Talafu West | 3.60 | 5.98 | 9.26 | 5.67 |
| Malitwang | 2.84 | 5.18 | 9.07 | 4.33 |
| Lanas II | 8.79 | 12.57 | 17.45 | 11.67 |
| Malibayong | 0.06 | 1.91 | 28.13 | 0.00 |
| Tangle | 3.71 | 6.23 | 9.82 | 5.67 |
| Total | 212.03 | 243.07<sup>*b*</sup> | 320.41 | 203.67 |

<sup>*a*</sup> Mean raw count per survey visit (summed across sex/age classes recorded on the same date, then averaged over the three – two for Nagbobong – survey dates per VP), not the total of all individual sightings summed across visits; the latter would count the same individuals repeatedly across repeat visits and is not on a comparable scale to the single-occasion, detection-corrected estimate at left. <sup>*b*</sup> Posterior median of the summed draws (each posterior draw’s abundance summed across all 18 VPs before taking the median), not the sum of the per-VP medians listed above; the two need not match because the median is not additive.

Our updated estimate of tamaraw density of 9.2 individuals km^−2^ (mean abundance across 2022–2026, with sampling variance removed) appears more biologically realistic, even if it ranks among the highest values we compiled from the literature for bovids of similar size (Table 1). If the population density of tamaraw was as high as 24 tamaraw km^−2^, we could expect strong density-dependence effects on demographic and annual growth rates, which would be visible as an especially low recruitment rate (Bonenfant et al. 2009). The fact that we have documented recruitment of young tamaraw during the most recent annual counts (years 2022–2024) (Bonenfant et al. 2023a) suggests that current density-dependence effects are moderate. It is possible that the annual burning of > 30% of the grassland in the CZM contributes to high quality forage at a critical timing for the species (Mentis & Tainton 1984; Thapa et al. 2022).

Because traditional burning occurs during the dry season, when plants have low palatability and moisture content, the CZM could support unnaturally high densities of tamaraw thanks to post-burn regrowth of the vegetation. It is possible that the carrying capacity of the study site might have been kept high because of this fire regime, therefore minimizing density-dependent feedback on population growth rate.

### Causes of the historical overestimation of tamaraw abundance

With a mean difference of almost 50%, the comparison of the tamaraw abundance at Mounts Iglit-Baco between historical counts and the double-observer estimator raises a conundrum. How can historical counts lead to abundance overestimation when we have evidence that detection probability < 1? Cases of overestimation of animal abundance are scarce in the literature, but the examples that we do have indicate that the underlying causes are diverse (Bonenfant et al. 2023b). We hypothesize that tamaraw identification errors accumulate with session number because of the way multiple-counts of the same individuals are dealt with during the consolidation phase. Because the tamaraw is a protected species, individuals cannot be marked, and during the consolidation phase of the historical count rangers are asked to remove individuals that may have been double-counted. The criteria for removing animals follow rules based on direction, time of observation, and group composition. If observers at neighboring VPs believe they saw the same individuals or group, then those tamaraw are removed from the total number of tamaraw seen that day. The same process applies to VPs across different sessions. As the time gap between sessions increases, it becomes less likely that group composition remains stable enough to confidently identify individuals. If a group splits while moving, its members may be counted more than once.

In addition, the proportion of animals removed during data consolidation — due to repeated observations of the presumed same individuals — varied over time, even though the observation methods remained consistent (see Supplementary Material 1). This suggests that the consolidation process may be influenced by the growing tamaraw population or potential changes in consolidation criteria used by rangers. As more individuals are observed each year, it becomes increasingly difficult for observers to determine whether a group has already been counted, likely resulting in fewer duplicates being identified in earlier years and contributing to underestimation in historical abundance estimates.

### Potential biases in the double-observer estimates

We emphasize that the double-observer estimator could still be biased and may slightly underestimate the tamaraw population size in Mounts Iglit-Baco Natural Park. There are several possible reasons for this. First, the model assumes complete independence between the two observers (Nichols et al. 2000), which was likely to be violated in some situations despite the intense training given to the field teams. Consequently, there may have been an overestimation of observer detection rates, leading to an underestimation of population size. A second unforeseen event that could skew our estimate is the unpredictable presence of military operations within the park during the 2023 count. That year, both the historical count (2 sessions cancelled) and the double-observer estimate dropped, which in turn made the following year’s growth rate biologically unrealistic (see Fig. 1a). We suggest that animals moved out of the CZM, or stayed hidden in forest patches, in response to the disturbance, and that this likely accounts for the abundance drop rather than a genuine population decline. These human activities almost certainly disturbed the animals (Andersen et al. 1996; Stephenson et al. 1996; Krausman et al. 2004) and may have led to unexpected tamaraw movements, including causing animals to venture outside of the CZM. This would mean that some individuals were not available for detection with our sampling design, and therefore give an underestimate of the total population (Péron & Garel 2019). Excluding year 2023 from the state-space model shifted the mean abundance estimate upward to *N̂* ^∗^ = _196.8_ 223.8 _253.6_, while leaving the mean growth rate basically unchanged (*r̄* = _−17.8%_ +6.7% _+45.1%_, *P*(*r* > 0) = 0.71). Because we carried out historical counts before the double-observer sampling, the induced human disturbance may reduce availability of tamaraw and hence, keep this abundance estimate low.

Finally, cloudy or rainy weather during some of the count sessions reduced visibility and may have lowered detection rates. Although weather was included as a covariate on detection probability in our models, these factors could still have biased the model results. For example, poor visibility and military activity could explain why the model predicted few tamaraw at the Magawang vantage point in 2022 (*N̂* = _14.0_ 18.4 _23.8_), despite the fact that this location showed the highest number of observed animals during the traditional count a week earlier (*C*_2022,Magawang_ = 99). Altogether, we believe that several factors may have negatively influenced the detection rates of observers, some of which could be accounted for in the models, and others which were not.

### Implications for tamaraw conservation

The Tamaraw Conservation Program Office (TCPO) has three main objectives (DENR 2019): (1) secure the short- and long-term viability of the tamaraw, (2) support the expansion of the current population within Mounts Iglit-Baco Natural Park, and (3) if possible, support the expansion of the tamaraw to other parts of Mindoro. The lower population of tamaraw in Mounts Iglit-Baco Natural Park that we report here will likely mean that conservation stakeholders will need to re-evaluate conservation actions to meet these objectives.

First, the small population size estimate provided by our models means that the fate of Mounts Iglit-Baco Natural Park tamaraw population is not as safe as previously believed. Smaller populations are at higher risk of demographic stochasticity and have lower genetic diversity, which in turn impacts population viability (Boyce 1992). Population and Habitat Viability Analysis (PHVA) models conducted in 2018 using the historical population size estimate (400-500) indicated that the Mounts Iglit-Baco Natural Park population was, in the absence of high levels of poaching, expected to persist at a stable equilibrium for the near future (DENR 2019). Unfortunately, in light of the new population estimates, it now seems that these conclusions are overoptimistic. We recommend that the PHVA models for the Mounts Iglit-Baco Natural Park population be re-run as a matter of urgency to better assess the viability of the population.

Second, the lower population size for the Mounts Iglit-Baco Natural Park population may also have broader implications for conservation plans for the species across Mindoro. At least two of the four other known tamaraw populations – in Mt. Calavite Wildlife Sanctuary, and Aruyan Malate – are extremely small and likely nonviable (Long et al. 2018). Current conservation plans envision exploring the translocation of animals from the Mounts Iglit-Baco Natural Park population to other sites to increase population size and genetic diversity, and therefore their long-term viability (DENR 2019). With approx. 200 individuals in the MIBNP population, such a scenario would be much more challenging than originally envisioned. Similar conclusions could be drawn regarding the removal of animals from the CZM to establish an ex-situ program. In both cases, removing a substantial number of animals from the CZM population could jeopardize its viability. One option to create a new population, as outlined in DENR (2019), would be to create conditions to allow natural expansion of the Mounts Iglit-Baco Natural Park population. To do this, it would be necessary to create a safe zone corridor that allows natural dispersal of the population from the core area in Mounts Iglit-Baco Natural Park to surrounding suitable areas. Considerable challenges remain in implementing these actions, though this conservation strategy is currently under discussion with relevant stakeholders.

## Conclusion

We emphasize that the overestimation of abundance in conservation could be more prevalent than anticipated. First, overestimated wildlife abundance is reported far less frequently than underestimation in the literature, which gives an inaccurate sense of a rather rare and unlikely situation (see Wessling & Surbeck 2023, for empirical cases on mammals). Second, observers might more easily accept a failure to detect an animal than counting the same individual several times within or between count sessions. Third, statistical methods for estimating population abundance that accommodate for multiple counts of individuals are scarce (see Table 1 for an overview), difficult to implement in the field, and therefore, rarely applied. Finally, the risk of overestimating population abundance can be more insidious than the risk of underestimating it, so it is easier to overlook. It is possible that lower population size estimates could be seen by authorities and conservation managers as indicative of unsuccessful conservation policies, which could lead to budget restrictions and/or loss of motivation for the project. In the case of the tamaraw, we stress that it is more likely that the overestimation has occurred since the start of the historical population count surveys, and that the magnitude of this overestimation has likely changed with abundance to an unknown extent. In practice, even if the long-term growth might have been overestimated because biases may co-vary with population abundance, it nevertheless highlights the effectiveness of conservation efforts and the usefulness of the historical tamaraw counts on Mindoro.

## ACKNOWLEDGMENTS

We also acknowledge the invaluable help of all rangers, students and volunteers who contributed to the annual tamaraw counts for two decades. Thanks are extended to Jean-Michel Gaillard for his critical and constructive reading of the manuscript.

## SUPPLEMENTARY MATERIAL

### S1: Inspection of historical count data at Mounts Iglit-Baco Natural Park

#### Statistical analyses of the historical abundance data

##### Results

We detected a significant (logistic regression of the consolidated fraction *f_t_* = *AN_t_*/*TN_t_* (with *TN_t_* the total number of tamaraw sighted and *AN_t_* the consolidated number retained after removing suspected repeat observations, in year *t*) on year, centered on 2004: logit(*f_t_*) = *β*_0_ + *β*_1_ × year + *β*_2_ × year^2^; *β*_0_ = −0.189 (SE = 0.057); *β*_1_ = −0.055 (SE = 0.010), *P* < 0.001; *β*_2_ = 0.0030 (SE = 0.0004), *P* < 0.001) non-linear pattern in the consolidated fraction over time (Fig. 1b, Fig. S1): a decline through the mid-2010s followed by a mild recovery in the most recent years (predicted consolidated fraction 45.3% at the start of the series in 2004 vs. 51.0% at the end in 2026).

**Figure S1.**
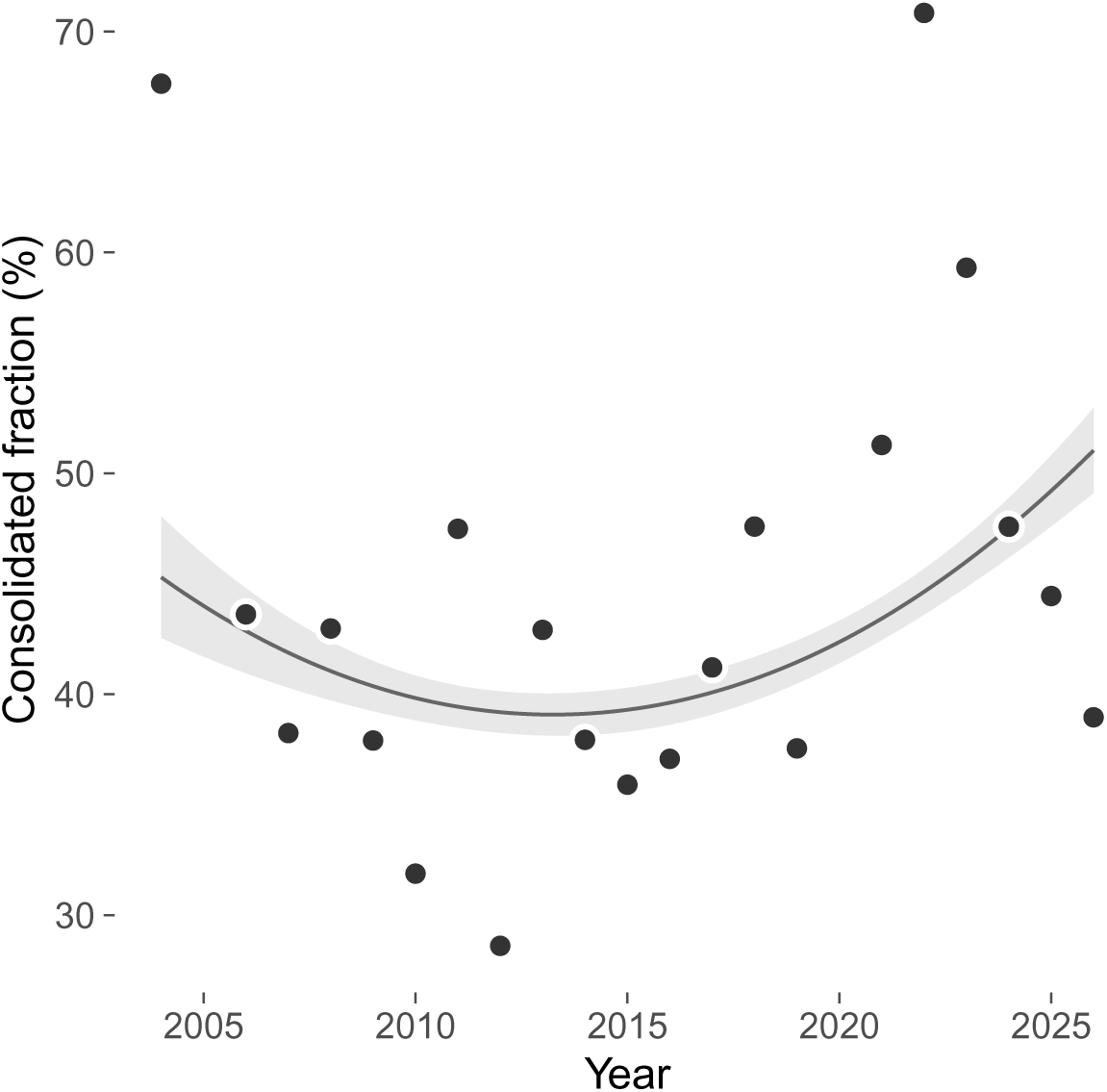
Consolidated fraction of tamaraw counts ( *f_t_* = *AN_t_*/*TN_t_*, %) by year, with the fitted quadratic logistic trend (line) and its 95% confidence band (shaded), corresponding to the regression reported above.

**Figure S2.**
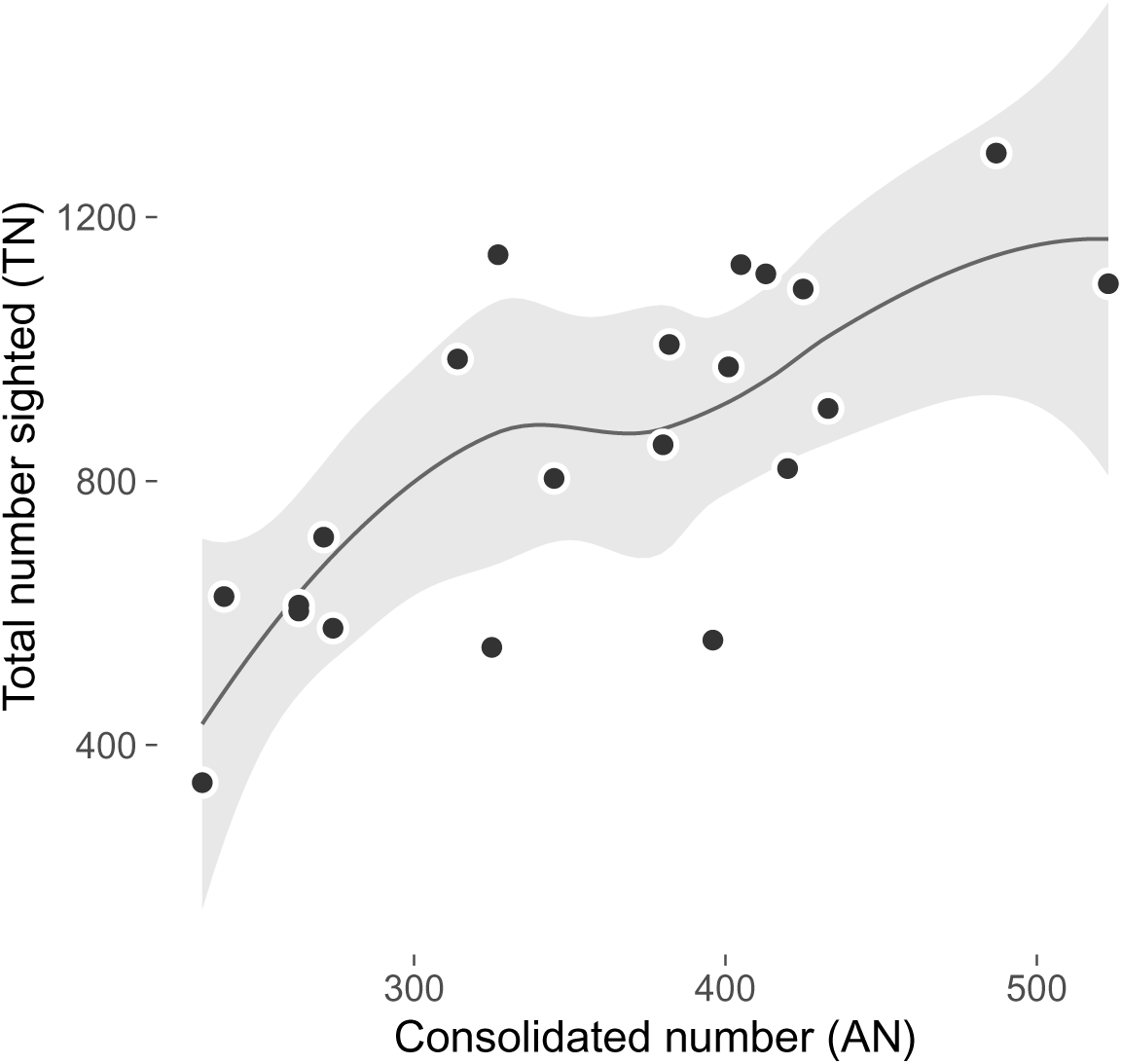
Total number of tamaraw sighted (*TN_t_*) versus the consolidated number (*AN_t_*), one point per survey year, with a loess trend line.

### S2: Spatial (Gaussian process) model exploration

We tested whether adding spatial structure to the VP-level random intercept on abundance improved the spatially-unstructured (independent and identically distributed, iid) version of the random effect reported in the main text. We present the geostatistical Gaussian process (GP) formulation in full detail below; two further conditional autoregressive parameterisations (a *k*-nearest-neighbour graph and a fully-connected distance-weighted graph) were also tested and performed similarly or worse (WAIC ∼1178 for the best-converged fit, against 1092.3 for iid and ∼1109 for the GP below) but are not detailed further here.

The GP model replaced the iid term *u_λ_*_,*s*(*i*)_ in the abundance submodel (Methods) with a spatially-correlated random effect *u_s_*, giving a proper multivariate normal prior with exponential covariance:

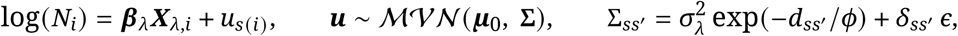

where *d_ss_*_′_ is the distance (km) between vantage points *s* and *s*^′^, *φ* a spatial range parameter estimated from the data (*φ* ∼ U(0.1, 3) km), *σ_λ_* the spatial standard deviation (*σ_λ_* ∼ U(0, 2)), and *δ_ss_*_′_ the Kronecker delta. As in the main abundance submodel, ***μ***_0_ was the historically-calibrated prior mean rather than a flat prior. We informed this parameter because the model failed to converge with a flat-prior at data-sparse VPs.

The nugget *ε* = 0.0001 is a small, fixed addition to the diagonal of **Σ**, a standard geostatistical practice to keep the covariance matrix numerically invertible, and not itself estimated. Without nuggets, **Σ** can become singular whenever two vantage points are located very close or at the same location. This configuration occurred for Loibfo I and Loibfo II for instance. The full JAGS code for the GP spatial model was:

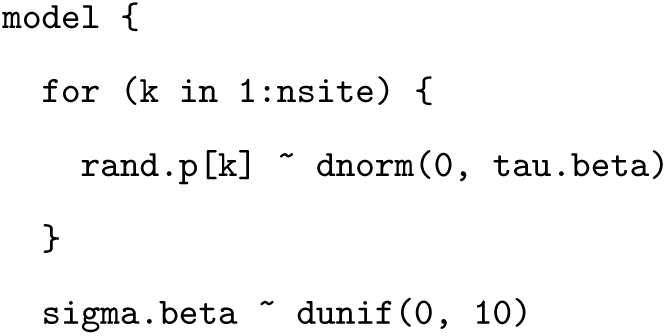
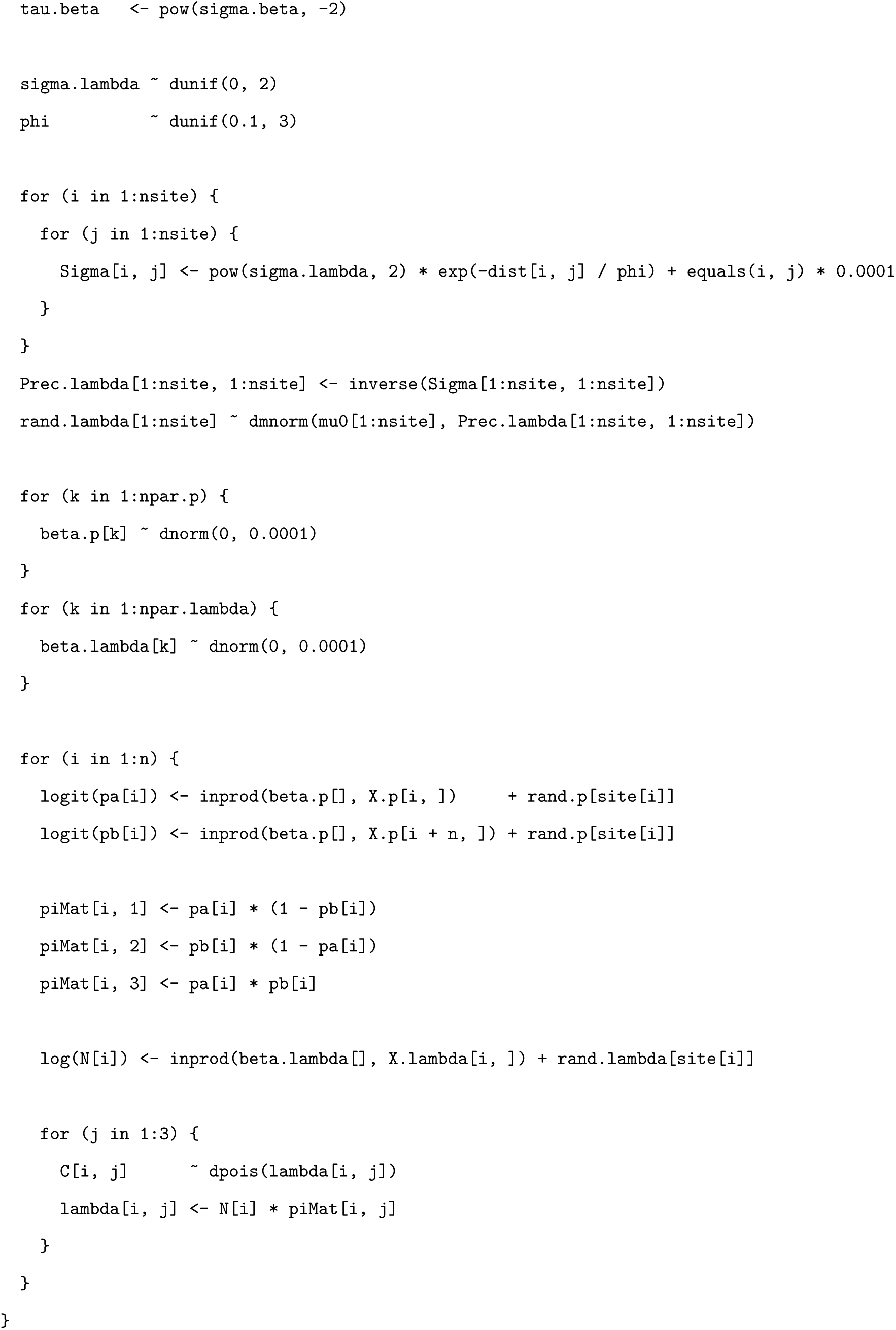

Here rand.lambda[s] is *u_s_*; dist the precomputed VP × VP distance matrix (km); mu0 the historically-calibrated prior mean vector; and X.lambda/X.p the abundance/detection design matrices (Methods). rand.p is *u_p_*_,*s*(_*_i_*_)_, sigma.beta is *σ_p_*, and beta.p/beta.lambda are ***β****_p_*/***β****_λ_* as defined in the main text. The detection and count likelihood are identical to the model used to fit the iid random effect.

The model estimated *φ̂* = 2.14 km (95% CI [0.53, 2.96]), suggesting that spatial correlation in local abundance decays substantially before reaching the typical distance separating vantage points (median pairwise distance 2.4 km; Fig. S4). WAIC did not improve on the iid model (1092.3) at ∼1109. We therefore retained the simpler iid random intercept for all analyses reported in the main text.

**Figure S3.**
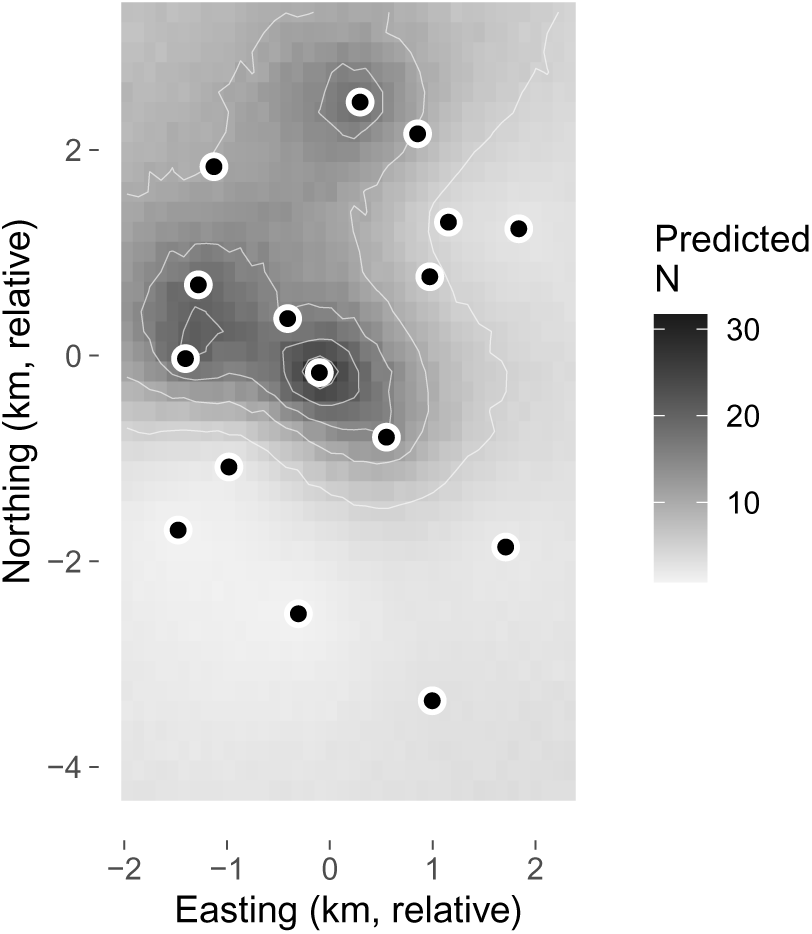
Predicted tamaraw abundance across the core zone of monitoring, kriged from the geostatistical (Gaussian process) spatial model’s posterior, with observed vantage points overlaid (dots).

**Figure S4.**
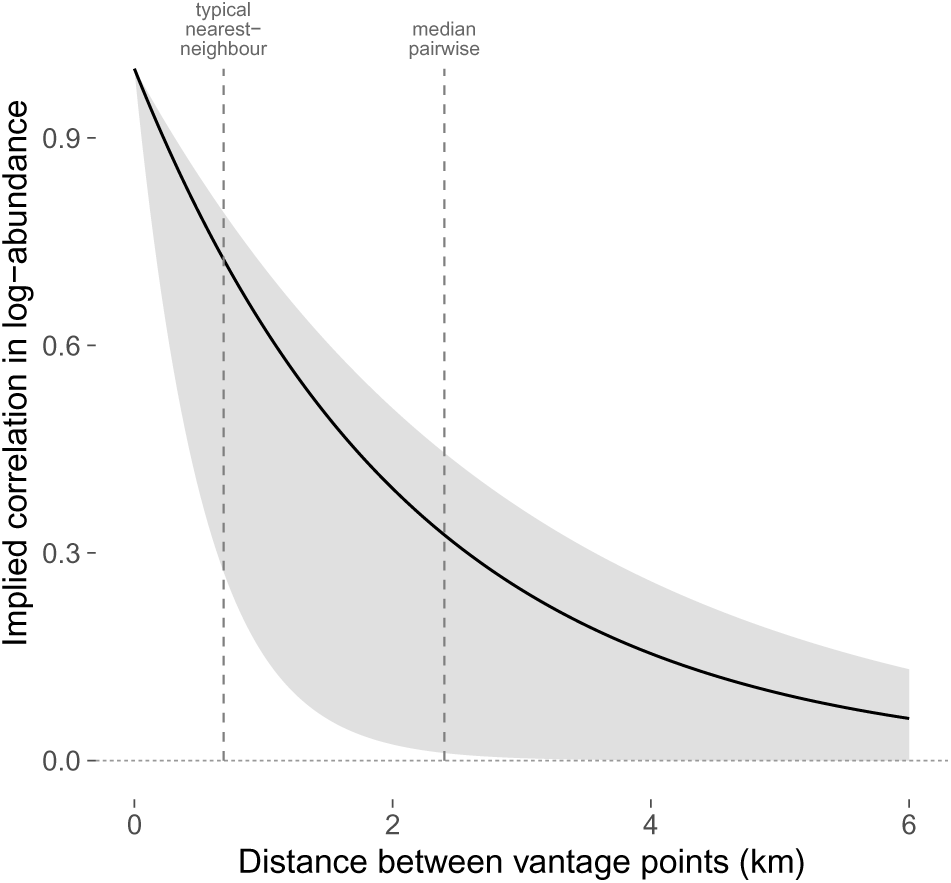
Implied correlation in log-abundance between two vantage points as a function of the distance separating them, derived from the geostatistical model’s posterior range parameter *φ̂* (median and 95% credible band). Dashed reference lines mark the typical nearest-neighbour distance and the median pairwise distance among the 17 vantage points used in this exploration.

